# *In situ* Nuclear Matrix preparation in *Drosophila melanogaster* and its use in studying the components of nuclear architecture

**DOI:** 10.1101/2021.09.30.462611

**Authors:** Rashmi U Pathak, Rahul Sureka, Ashish Bihani, Parul Varma, Rakesh K Mishra

**Affiliations:** Centre for Cellular and Molecular Biology, Uppal Road, Hyderabad-500 007, India; EMBL, Rome, Italy; Department of Neuroscience, Development and Regenerative Biology, The University of Texas at San Antonio, Texas, USA

**Keywords:** Nuclear matrix, nuclear architecture, *in situ* NuMat

## Abstract

The study of Nuclear Matrix (NuMat) over the last 40 years has been limited to either isolated nuclei from tissues or cells grown in culture. Here, we provide a protocol for NuMat preparation in intact *Drosophila melanogaster* embryos and its use in dissecting the components of nuclear architecture. The protocol does not require isolation of nuclei and therefore maintains the three-dimensional milieu of an intact embryo, which is biologically more relevant compared to cells in culture. One of the advantages of this protocol is that only a small number of embryos are required. The protocol can be extended to larval tissues like salivary glands and imaginal discs with little modification. Taken together, it becomes possible to carry out such studies in parallel to genetic experiments using mutant and transgenic flies. This protocol, therefore, opens the powerful field of fly genetics to cell biology in the study of nuclear architecture.

**Summary:** Nuclear Matrix is a biochemically defined entity and a basic component of the nuclear architecture. Here we present a protocol to isolate and visualize Nuclear Matrix *in situ* in the intact embryos and tissues of *Drosophila melanogaster* and its potential applications.

## Introduction

It is well established that nucleus is compartmentalized, and its functional domains are dynamically linked. Nuclear Matrix (NuMat) is a structural framework involved in organization of internal nuclear architecture. NuMat was first described by Berezney and Coffey in 1974 as a nuclear sub-structure consisting of a meshwork of ribonucleo-proteinaceous filaments which resists extraction by non-ionic detergents and high salt concentrations. The NuMat persists even after the chromatin has been completely removed^1^. The discovery of NuMat is followed by several studies where it has been proposed as a substratum on which various nuclear processes such as replication, transcription, DNA repair, splicing and chromatin remodelling can happen. It is shown that NuMat association of the components involved in these processes indeed facilitates these nuclear processes^2–6^. Biological relevance of NuMat has been extensively reviewed over the years^7–9^. However, most of the studies on NuMat have been carried out either in cultured cells or isolated nuclei, both of which provide valuable insights but have the caveat of disturbing the nuclear architecture and genome organization and hence may not reflect the *in vivo* conditions faithfully.

Till now, it has been challenging to harness the power of genetics in the study of NuMat because of the lack of a methodology to visualize NuMat in the context of the intact organism or tissue. Here, we describe a method to prepare NuMat *in situ* in the developing embryo and larval tissues of *D. melanogaster.* Our method bridges this gap as it makes it possible to visualize NuMat *in situ* in the organism. *D. melanogaster* is one of the preferred model organism for variety of reasons, including the sheer abundance of available mutants. *Drosophila* genetics combined with the presented method of *in situ* NuMat preparation has the potential to provide a robust method to study various components of nuclear architecture.

## Results

### *In situ* NuMat preparation retains the characteristic features of nuclear architecture

The main experimental steps of *in situ* NuMat preparation have been outlined in the form of a workflow in Figure 1. In an intact nucleus, the NuMat is concealed by dense chromatin mass which is removed by extraction with non-ionic detergent and salt followed by DNase I digestion. The NuMat thus revealed, consists of a nuclear lamina, an internal matrix composed of thick polymorphic fibres and ribonucleoprotein particles and remnants of nucleoli. To assess the quality of nuclear matrices prepared by the *in situ* NuMat preparation method, we visualized it by TEM and confocal imaging (Figure 2).

**Figure 1:**
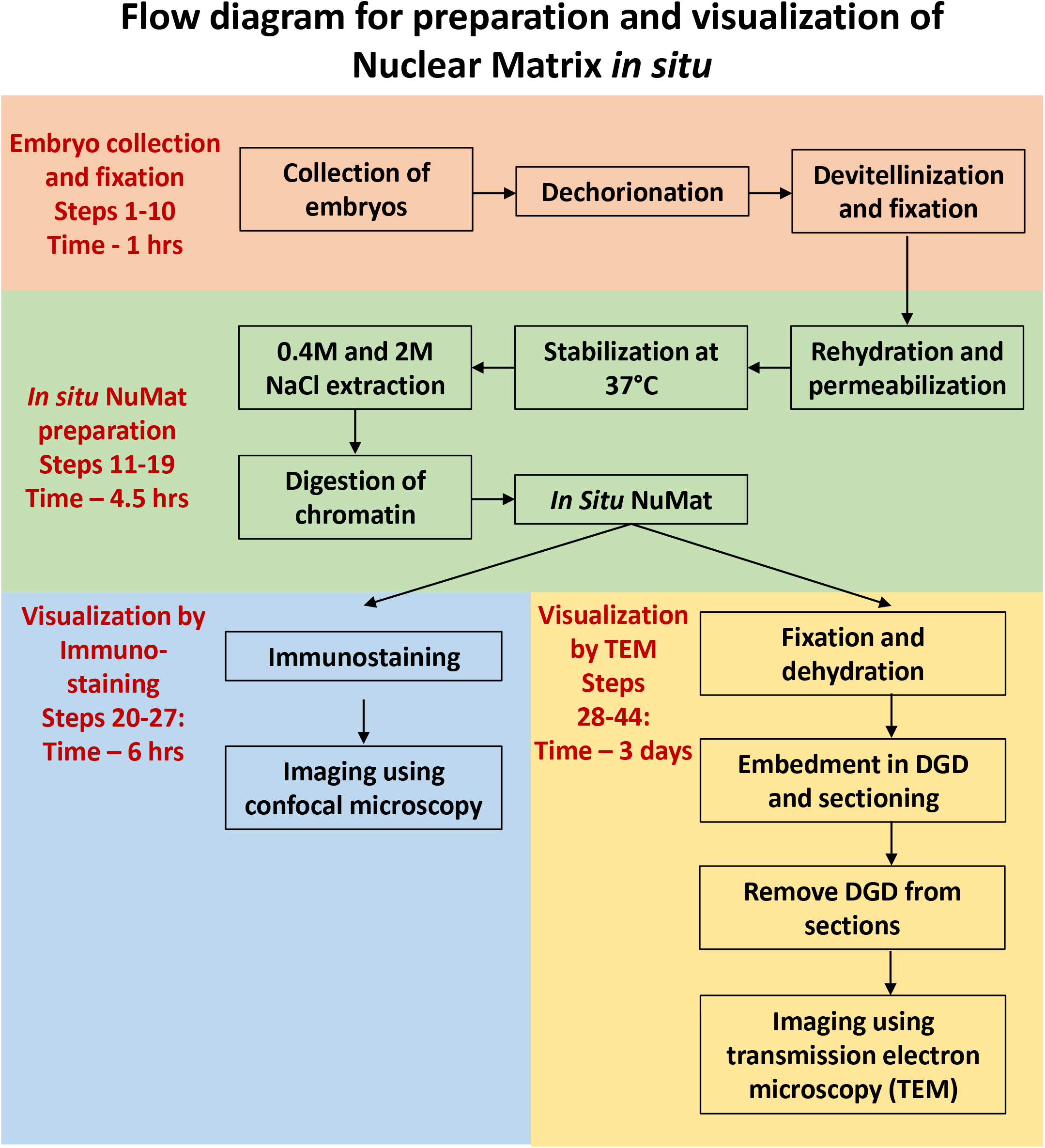
The scheme of steps for *in situ* NuMat preparation.

**Figure 2:**
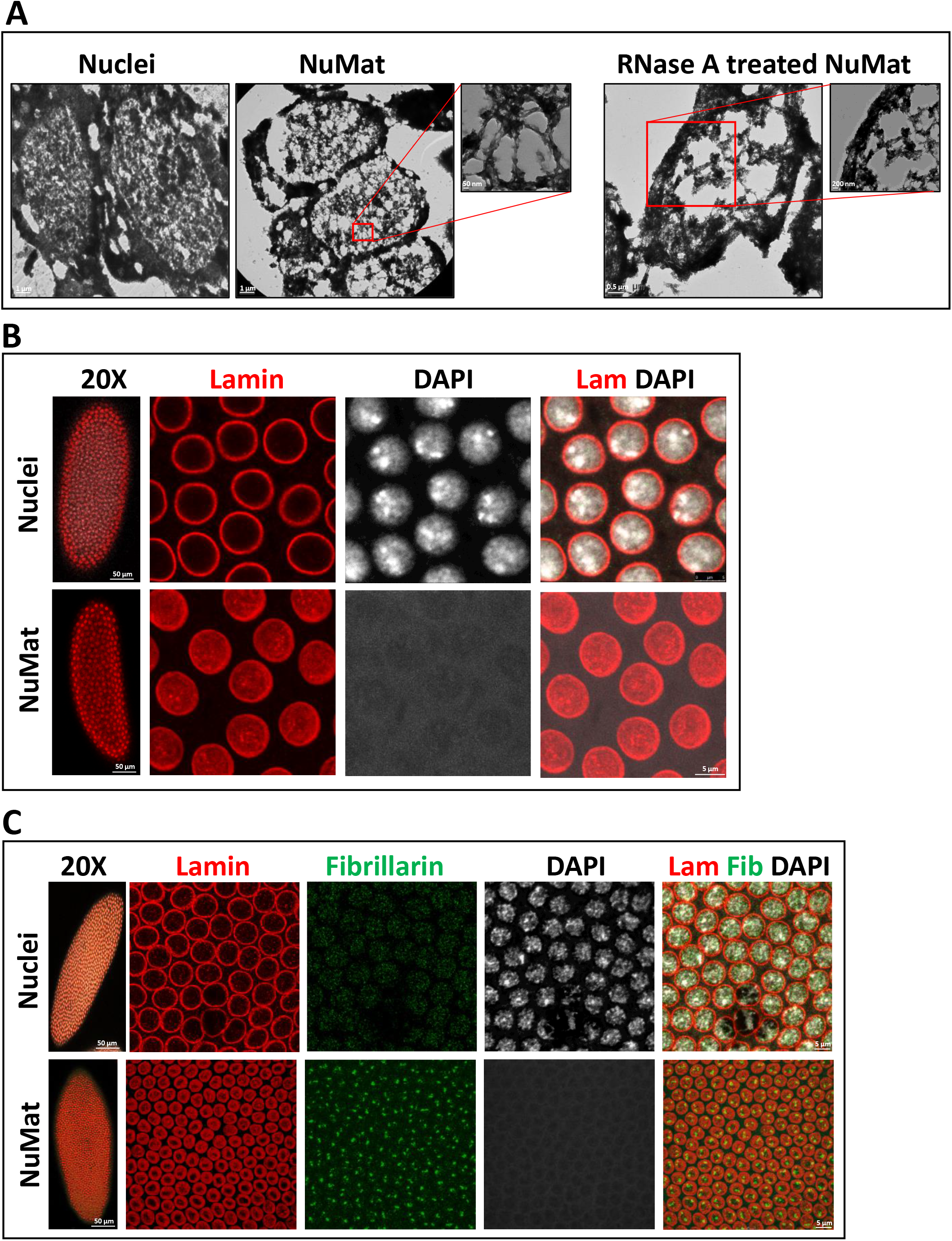
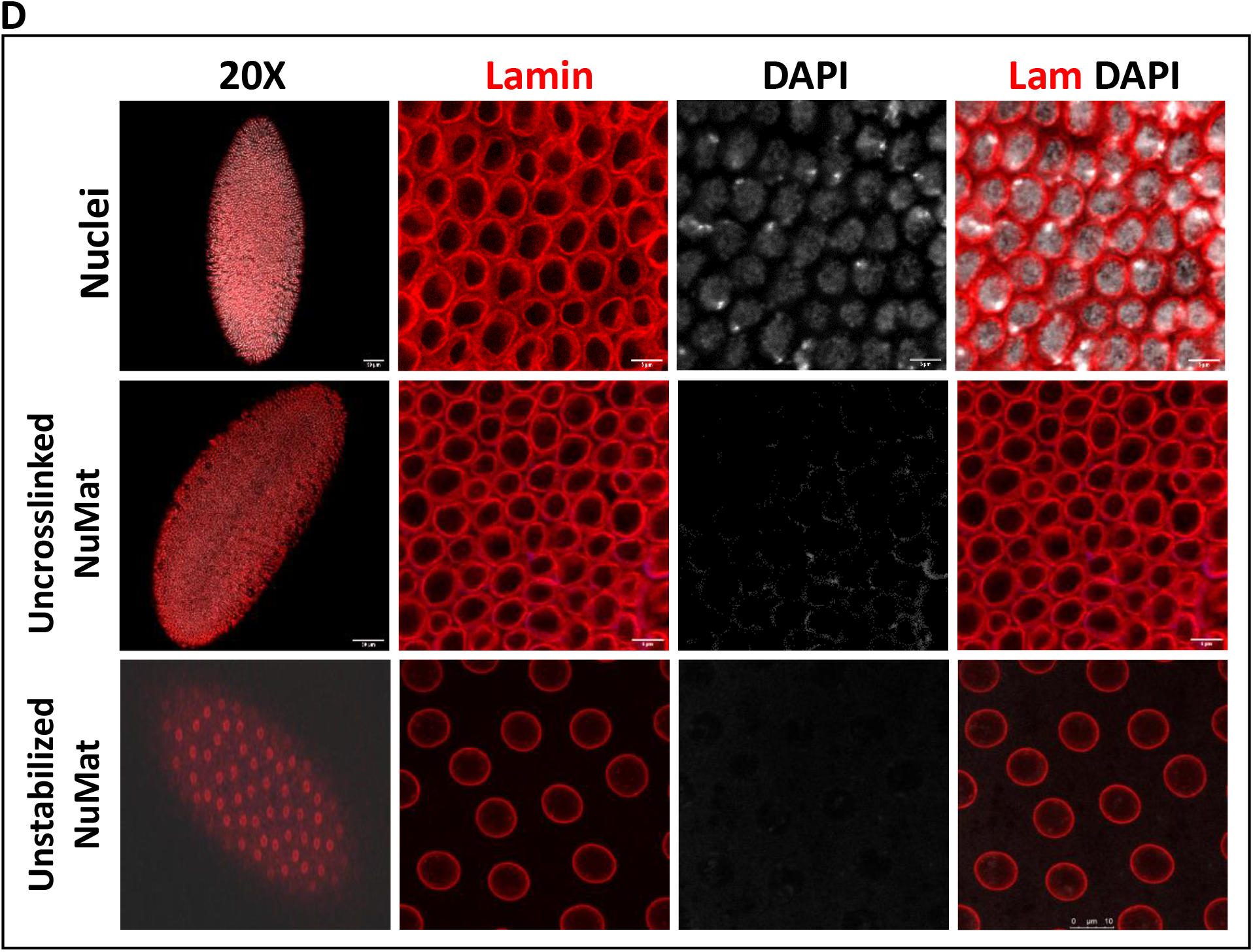
Visualization of *in situ* NuMat. *D. melanogaster* embryos at early stages of development (0-2 hr) were used to prepare nuclear matrices *in situ.* **A.** Visualization by TEM. Images obtained by TEM of resinless sections of embryos carrying intact nuclei, *in situ* NuMat and RNase A treated *in situ* NuMat. The fine filaments seen in NuMat, are lost upon RNase A treatment, leaving large gaps in the nuclear structure and leading to collapse of nuclei. **B.** Visualization by confocal microscopy. Unextracted embryo and embryo with *in situ* NuMat, were immuno-stained with anti-Lamin Dm0 and DAPI and imaged by confocal microscopy. In unextracted embryos, Lamin Dm0 appears as a ring at the nuclear periphery of intact nuclei. After *in situ* NuMat preparation, no DAPI staining is observed in the nucleus, as chromatin has been digested and extracted out. Lamin Dm0 staining can now be seen in the nuclear interior as well. **C.** Remnants of nucleolus remain associated with *in situ* NuMat. Confocal images of unextracted embryo and embryo with *in situ* NuMat, were immuno-stained with anti-Fibrillarin, anti-Lamin Dm0 and DAPI. Loss of DAPI staining indicates extraction of chromatin. *In situ* NuMat shows prominent staining with fibrillarin indicating remnants of nucleolus remain associated with the nuclear substructure. **D.** *In situ* NuMat prepared without crosslinking or stabilization. *In situ* NuMat is efficiently prepared as evident by absence of DAPI staining, even when the embryos are not crosslinked or are not stabilized. The circular morphology of the nuclei remains intact, but internal lamin staining is not visible.

Early studies have shown that the filamentous network in an extracted nucleus is best illustrated when embedment free sections are visualised by TEM. The conventional epoxy embedded sections can obscure important biological structures, thus resinless sections have been routinely used to visualize NuMat. We used resinless sectioning followed by TEM imaging to assess the ultrastructure of *in situ* NuMat prepared in early *Drosophila* embryos (Figure 2A). In line with previous reports, the intact nucleus is filled with a dense network of chromatin and soluble proteins. After extraction with detergent, salt and DNase I, the nuclear interior is visualised as a network of filaments bound by lamina. Treatment of NuMat with RNase A results in loss of the internal fibres and aggregation of ribonucleoprotein particles, highlighting the importance of RNA in matrix organization. The RNA-containing NuMat appears as a self-supporting three-dimensional structure which collapses after removal of RNA. The RNA depleted NuMat is also markedly distorted in overall shape. These observations suggest that the nuclear matrices prepared *in situ* in intact *Drosophila* embryos are ultra-structurally similar to NuMat prepared in cultured cells by traditional methods.

The chromatin depleted NuMat is a core structure where lamins are retained along with a unique set of nuclear non-histone proteins that resist salt and detergent extraction. To visualize these proteins, we immuno-stained the *in situ* NuMat with anti-Lamin Dm0 and anti-Fibrillarin antibodies (Figure 2B and 2C). We observe that Lamin Dm0 grossly defines the nuclear morphology. Apart from forming a meshwork adjacent to the inner nuclear membrane, it is also present in the nuclear interior. Further we also see that the well characterised structural feature of eukaryotic nucleus, the nucleoli, is prominently visible after *in situ* NuMat preparation (Figure 2C). The complete removal of chromatin (absence of DAPI staining) confirms the efficiency of extractions.

Previous studies on NuMat preparations on cell lines have shown that formaldehyde treatment is essential for TEM visualization of the characteristic fibrillar meshwork of NuMat. However, it is not known if formaldehyde crosslinking and stabilization is essential for *in situ* NuMat preparations in *Drosophila* embryos as well. In order to access the necessity of these treatments, we excluded them from the protocol. As seen in Figure 2D, the overall shape and morphology of nuclei remains intact even if the fixation or stabilization steps are omitted from the NuMat preparation protocol. Complete removal of chromatin (no DAPI staining) indicates that NuMat is successfully prepared. However, internal lamin is not seen in uncrosslinked as well as unstabilized NuMat preparation. Thus, we observe that crosslinking and stabilization steps can be excluded from the protocol. However, based on Lamin Dm0 visualization, the *in situ* NuMat prepared after crosslinking and stabilization appears to be more intact.

### *In situ* NuMat preparation can be extended to late embryos and larval tissue

The strength of *in situ* protocol lies in the proposition that the power of *Drosophila* genetics can be harnessed to uncover novel molecular players with a role in nuclear architecture. However, in *Drosophila*, the effects of a genetic manipulation may not manifest in the nuclei of early embryos because of masking due to maternal deposition of the molecule of interest in the embryo. In such cases, the phenotype would be visible only at later stages of embryonic development. This necessitates that the protocol works effectively in older embryos or specific tissues in the larval or adult stages. As seen in Figure 3, the chromatin digestion and salt extraction, works well in older embryos (Figure 3A) as well as larval salivary glands (Figure 3B). Negligible staining with DAPI indicates that bulk of chromatin is effectively removed and Lamin Dm0 in the nuclear interior is revealed defining the NuMat.

**Figure 3:**
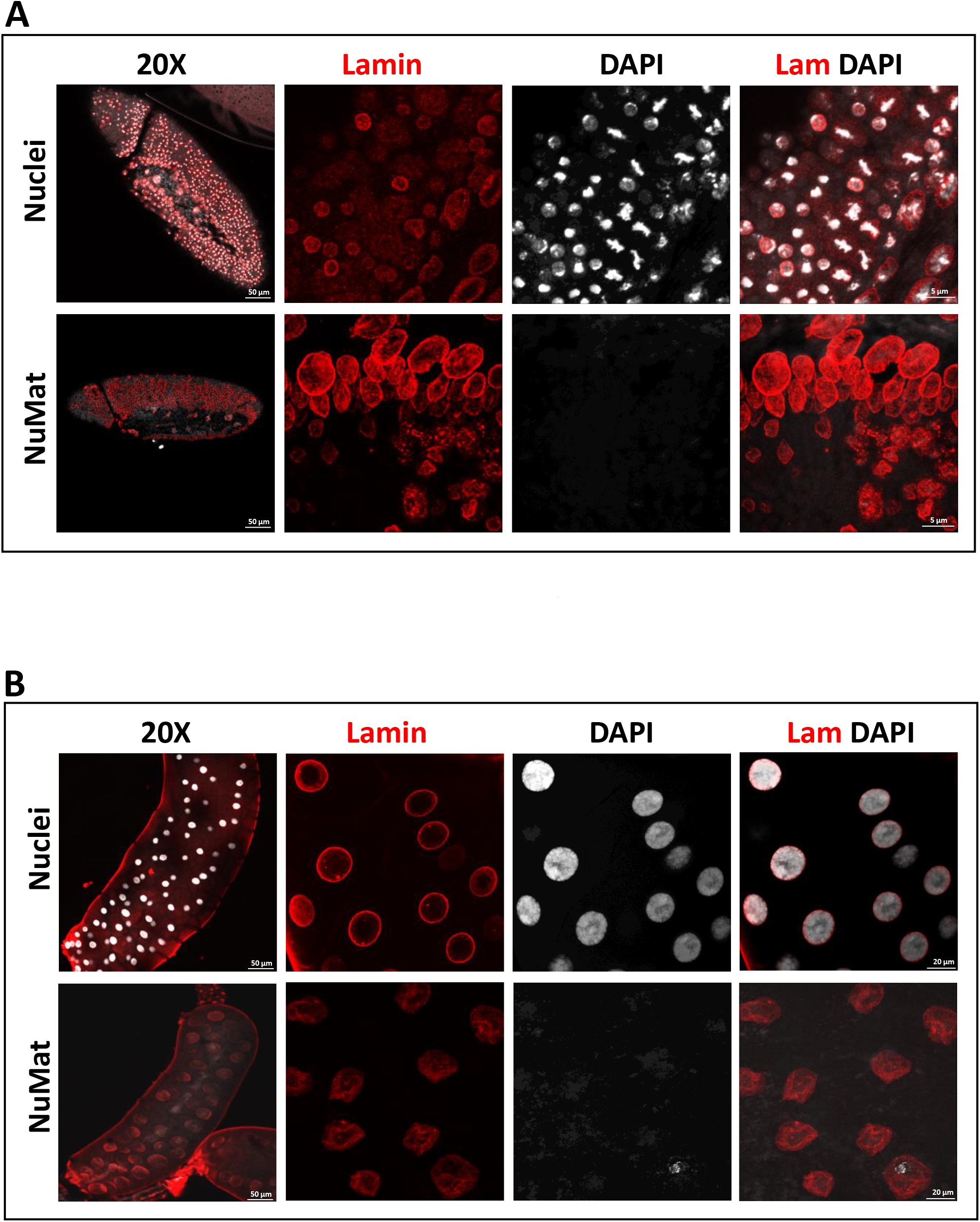
*In situ* NuMat preparation with late *D. melanogaster* embryos and larval tissues. **A.** *In situ* NuMat prepared with embryos at late stage of development shows that the digestion and extraction of chromatin (assessed by loss of DAPI staining) works well with different types and layers of cells present in a developing and differentiating embryo. Intra-nuclear Lamin Dm0 is revealed after the nuclear matrices are prepared. **B.** *In situ* NuMat prepared with *D. melanogaster* 3^rd^ instar larval salivary glands shows that the bulk of chromatin present in polytene chromosomes is efficiently extracted (assessed by loss of DAPI staining) to reveal the nuclear architecture of salivary gland nucleus.

### Use of *in situ* NuMat preparation to study dynamics of proteins during mitosis

The early *Drosophila* embryo is a treasure-trove of interesting biological phenomenon. The nuclei in the syncytial embryo undergo 13 rounds of division. The cell cycle lasts only for 8 min during the early stages and progressively slows down to 18 min for the cycle 13^10^. The rapid nuclear divisions at this stage increases the probability of catching an embryo with mitotic wave at the surface. It is possible to fix embryos during such a mitotic wave and study the cell cycle related dynamics of a nuclear constituent of interest.

Here we take an example of the protein BEAF 32 (Boundary Element Associated Factor), a known NuMat associated protein, to show how the association of this protein with nuclear architecture varies through cell cycle. BEAF 32 bound DNA boundary elements are known to tether to nuclear architecture by virtue of the interaction of the protein with NuMat^11^. *In situ* NuMat prepared on an early embryo that had a mitotic wave on display, shows that BEAF 32 remains associated with nuclear architecture at different stages of mitosis (Figure 4).

**Figure 4:**
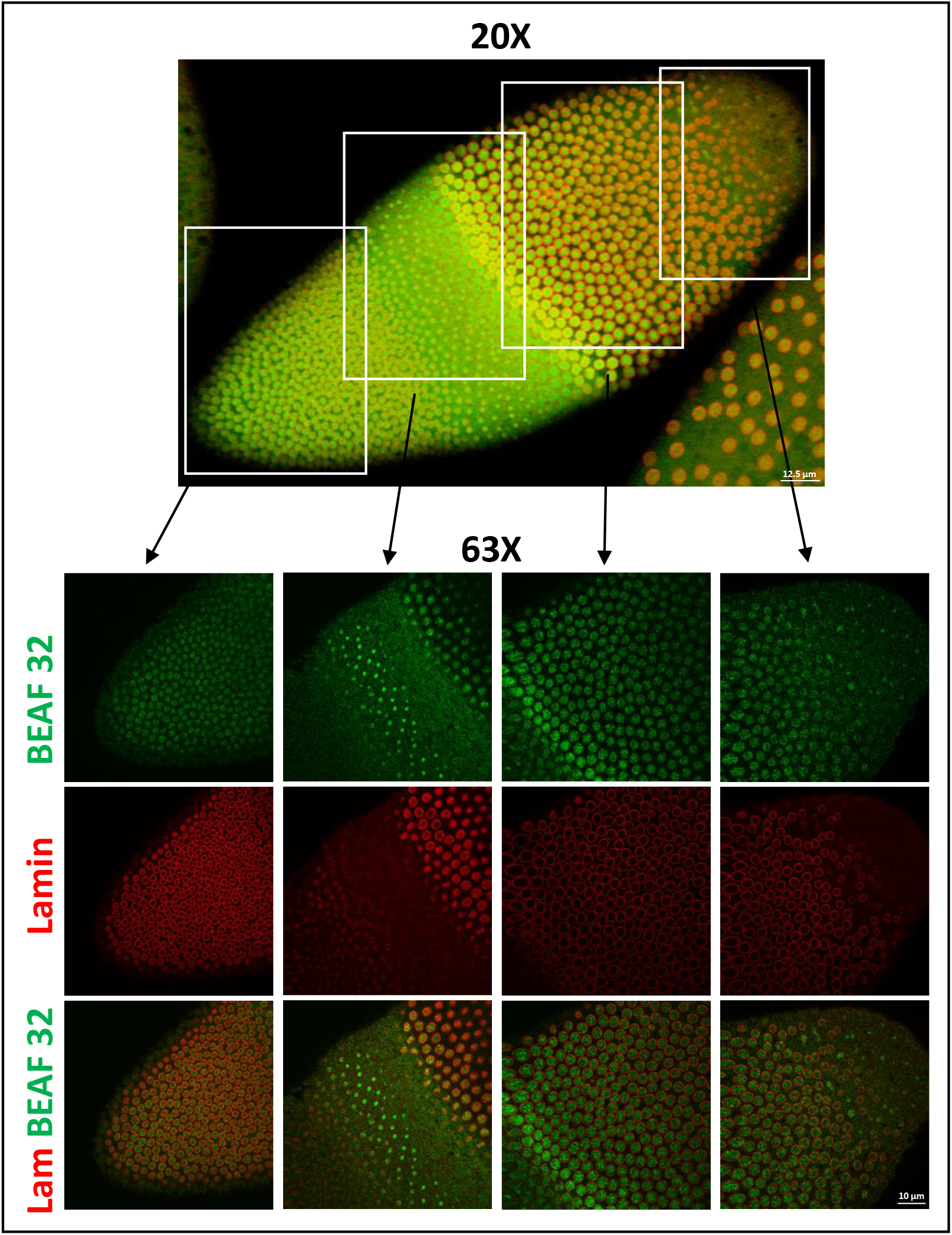
*In situ* NuMat preparation using early *D. melanogaster* embryo with a mitotic wave. *In situ* NuMat prepared with early syncytial embryos, captures a snapshot of an embryo with nuclei at different stages of mitosis. Immuno-staining with anti-Lamin Dm0 and anti-BEAF 32 reveals the dynamics of these nuclear proteins at different mitotic stages. A subset of BEAF 32 stays associated with mitotic nuclei even when the nuclear envelope (defined by Lamin Dm0) is dissolved.

### *In situ* NuMat preparation can be used in conjunction with fly genetics

One of the common strategies used to study the possible role of a candidate protein in nuclear structure and function is to immuno-stain the protein for its localization. Disruption/depletion of a particular protein by mutation/RNAi is also used as an effective tool for analysing gene function. *Drosophila* as a model organism is well suited for such studies. A vast repertoire of tagged-fly lines available publicly, is a valuable resource that makes it possible to study a novel protein, without worrying about the availability of its antibody. The BDGP gene disruption collection that disrupts ~40% of fly genes, provides a public resource that facilitates the application of *Drosophila* genetics to diverse biological problems^12 13^. On similar lines, the Transgenic RNAi Project (TRiP) has generated transgenic RNAi fly stocks that use Gal4/UAS system to induce RNAi silencing of specific genes. Embryos from such fly lines can be used for *in situ* NuMat preparation for architectural studies.

Here again, we take an example of differential association of isoforms of BEAF 32 protein with the nuclear architecture. BEAF 32 exists as two isoforms, 32A and 32B, that form a hetero-trimer to bind to the chromatin. The only antibody available for BEAF 32 (from DSHB) does not differentiate between the isoforms. We, in our lab have generated two transgenic fly lines one of which had Myc tagged BEAF 32A transgene in it and the other had FLAG tagged BEAF 32B transgene present in it. Crossing them as shown in Figure 5A, gives flies with Myc tagged BEAF 32A on chromosome 2 and FLAG tagged BEAF 32B on chromosome 3 simultaneously, in the F1 generation. Such a fly helped us to study both the isoforms of BEAF 32 in a single nucleus. Salivary glands from 3^rd^ instar larvae from this fly line were stained with anti-Myc and anti-FLAG antibodies simultaneously. As seen in figure 5B, BEAF 32A and 32B colocalize on most of the sites on the polytene chromosome in an intact nucleus. However, after *in situ* NuMat preparation, BEAF 32A staining is reduced to negligible, while BEAF 32B is retained predominantly in the NuMat. This experiment shows that the two isoforms of BEAF 32 interact differently with the underlying nuclear architecture.

**Figure 5:**
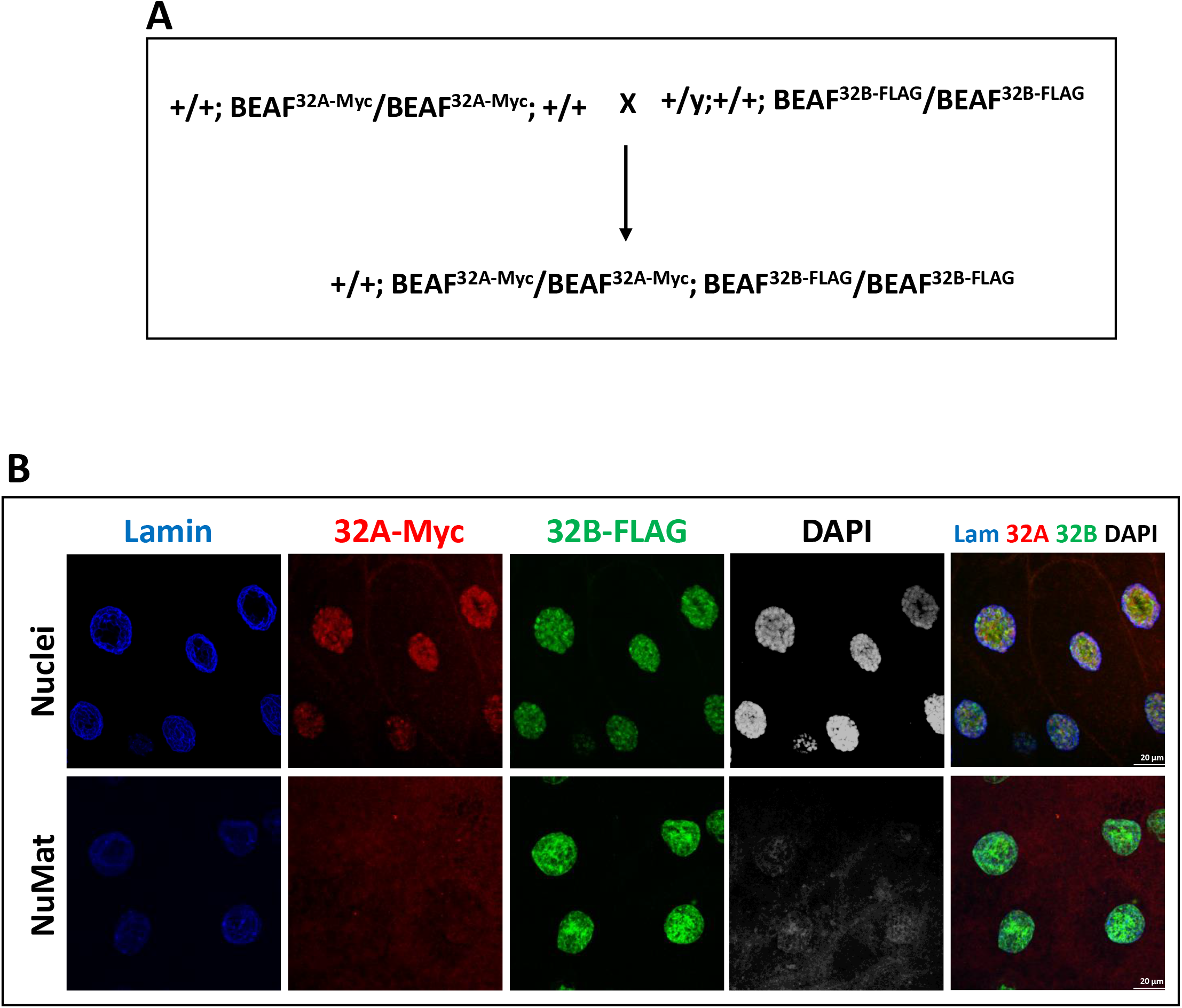
*In situ* NuMat preparation in conjunction with mutant fly. **A.** Fly cross scheme to generate a fly line carrying tagged isoforms of BEAF 32 in the same fly. **B.** Immuno-staining of unextracted/*in situ* NuMat prepared salivary glands with anti-Lamin Dm0, anti-Myc and anti-FLAG antibodies. Myc-tagged BEAF 32A and FLAG-tagged 32B, colocalize on several bands of the polytene chromosome in the salivary gland nuclei. After *in situ* NuMat preparation, 32A gets extracted out and 32B remains associated with NuMat.

### *In situ* NuMat preparation can be used to unravel novel components of nuclear architecture

The *in situ* NuMat preparation can be utilized to understand the composition and dynamics of nuclear architecture. We demonstrate this by taking an example of staining for polymeric actin in the nucleus (Figure 6). Presence of actin in the nucleus is debated as it is still a matter of investigation that in what form, and what for, the protein is present in the nucleus. For a while it was believed that nuclear actin is predominantly monomeric (globular, G-actin). This was partly due to inability to visualise F-actin in the nucleus. However, recent research has made it possible to see nuclear actin filaments under the microscope. Since then, transient nuclear actin filaments have been described in several cellular processes such as serum response, DNA damage, cell spreading, chromatin de-condensation and gene transcription^14^. Our *in situ* NuMat preparation provides a simple way to visualise polymeric actin in the nucleus, that otherwise remained concealed under the bulk of chromatin. When we stained the NuMat preparation with GFP labelled phalloidin (highly selective for F-actin), we observe that polymeric actin is indeed present in nucleus. *In situ* NuMat prepared in early (Figure 6A) or late (Figure 6B) stage embryo removes the chromatin to reveal polymeric actin in the nucleus.

**Figure 6:**
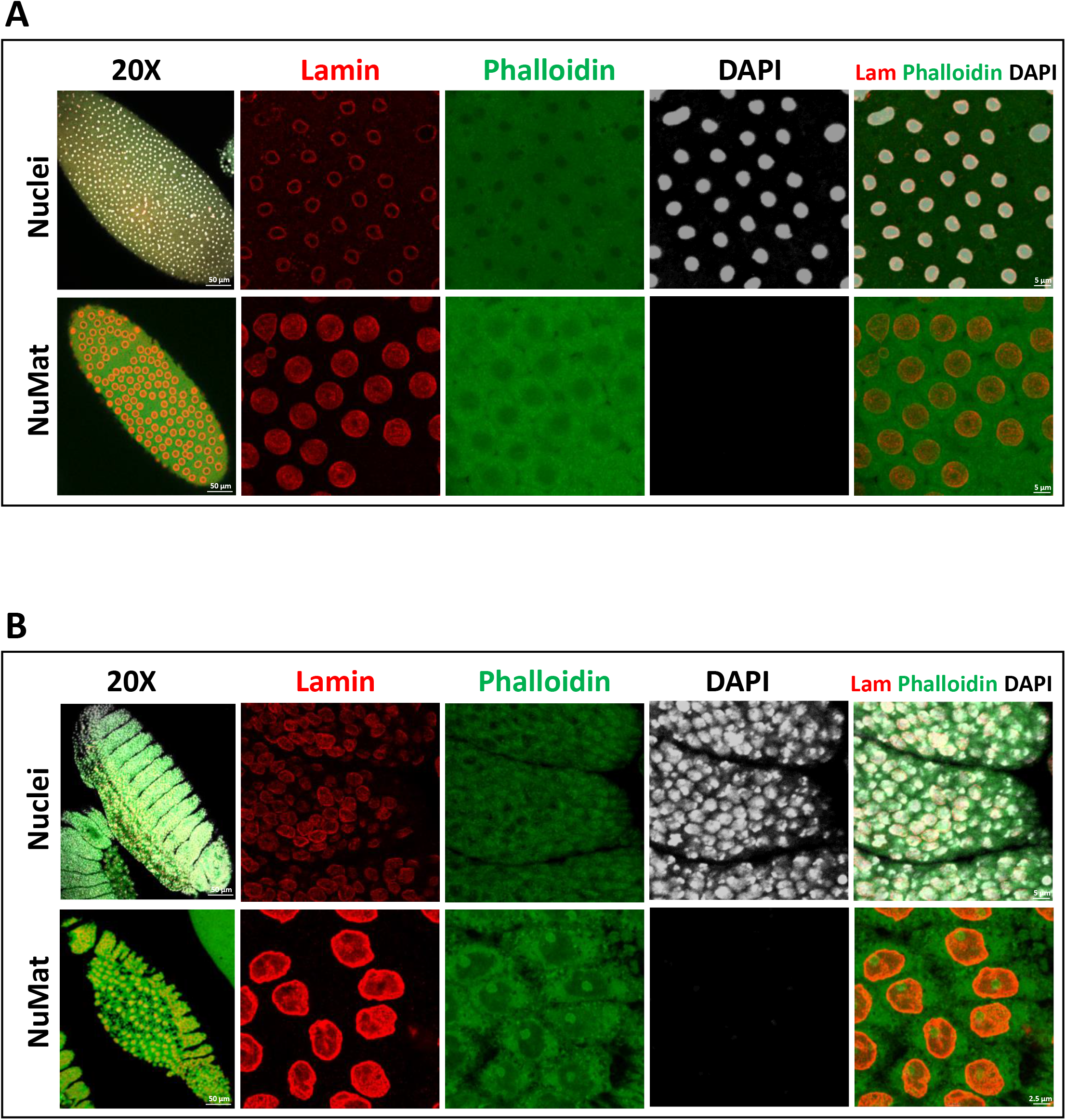
*In situ* NuMat preparation protocol to reveal polymeric actin in the nucleus. *Drosophila* embryos at early **(A)** or late **(B)** stage of development were stained with Phalloidin, Lamin Dm0 and DAPI. In control embryos, the polymeric actin is not visible in the nucleus as it remains obscured by chromatin. After *in situ* NuMat preparation, Phalloidin staining can be seen in the nucleus.

## Discussion

The formidable task of organizing almost a meter of DNA into a ~10uM diameter cell nucleus is facilitated by NuMat. It is proposed to be the framework that unifies the nuclear structure and function. This important nuclear sub-structure is evidently structurally quite complex. Composition of NuMat varies from cell to cell, far more than cytoplasmic or chromatin fraction, indicating its functional significance^15^. Biochemical extraction experiments and TEM imaging conducted through 1980s have defined the NuMat to be majorly ribo-proteinaceous in nature. These experiments also provide wealth of data regarding the variations observed in the NuMat, when choice of salts/detergents or sequence of extraction is altered. However, all studies invariably agree on one point where a distinct nucleoskeletal structure is observed upon the removal of bulk of chromatin. Further studies have linked NuMat functionally to all of the nuclear processes. The DNA needs to interact with the sites where nuclear functions are performed (in the NuMat) and this necessity forms the basis of another layer of regulation of chromatin function. This regulation of chromatin function by nuclear architecture defines the concept of ‘spatial epigenetics’^16^. However, the specific molecular players still remain unexplored. The NuMat as a nuclear substructure remains an enigma due to the limitations posed by the lack of a screening technique, where the queried component is removed/overexpressed and its effect on nuclear architecture is visualised. Limitations are further enhanced by the technicalities involved in imaging NuMat. Live imaging has not been possible, as most of the NuMat components remain obscured by chromatin. On the other hand, looking at an extracted isolated nucleus or cells in culture also have limited utility.

Here we present a method for visualizing structural components of the nucleus, in the background of an experimental screening setup. Our method for *in situ* NuMat preparation makes it possible to study nuclear architecture in conjunction with the vast genetic resources of *Drosophila*. It allows the observation of NuMat in the *in vivo* context of a developing *Drosophila* embryo which has been very challenging until now. It can thus be used to study the cell cycle related dynamics of the molecule of interest. The technique can also be used to study nuclear architecture in various other *Drosophila* tissues like the salivary glands, imaginal discs, etc. Once the *in situ* NuMat has been prepared, this technique is not only limited to immuno-staining and TEM but can also be used for DNA and RNA-FISH experiments. We have previously demonstrated such an application by studying the role of AAGAG RNA in nuclear architecture by using this technique^17^. As NuMat is the site of various nuclear functions like replication, transcription, DNA repair and splicing, this technique can be used to probe deeper into these questions in the nuclear architectural context in the embryo.

Our method should not be confused with a method developed previously by Capco et. al.^18^ in which whole cell mounts of 3T3 cell lines were extracted in buffers with physiological ionic strength without isolating nuclei. Their method has also been referred to as *in situ* NuMat in reports, as it extracts the whole cell to derive the NuMat and no nuclei purification step is involved^19^. Here we adopt the classical biochemical techniques for making NuMat and extend it to intact *Drosophila* embryos/tissues. This adoption enables the visualization of nuclear architecture, in the background of an experimental setup, and compliments biochemistry, cell biology with *Drosophila* genetics.

Our protocol for *in situ* NuMat preparation is simple and does not require advanced technical knowledge. It can be performed by any researcher with basic experience in molecular biology. Further comparison with other existing protocols for NuMat preparation illustrates that the *in situ* NuMat preparation has several advantages. For instance, most protocols of NuMat preparation either involve isolation of pure nuclei from tissues or use cell lines which have been grown in cell culture for a number of passages. Isolation of pure nuclei requires expertise, is time consuming and technically challenging, and often requires specialized equipment such as homogenizers and ultracentrifuges. On the other hand, cultured cells are often grown as 2D monolayers which does not entirely reflect *in vivo* condition because of the lack of 3D cues^20,21^. However, our protocol does not require isolation of pure nuclei which reduces the time of preparation, hence minimizing the possibility of adventitious crosslinking by sulfhydryl oxidation that has been reported for lengthier protocols^22^, and is easily adaptable. Moreover, our technique uses the whole developing embryo to prepare NuMat *in situ*, thereby causing minimal disturbance and thus reflects the *in vivo* conditions most faithfully. Taken together, we suggest that *in situ* NuMat preparation will facilitate in depth analysis of nuclear architecture in the context of nuclear functions. An important advantage of our protocol is the requirement of low amount of sample, and therefore can be used to answer biologically relevant questions with very few embryos obtained by a genetic screen, in contrast to the large amount of tissue, embryos or cells required for biochemical experiments^23–26^. As this technique uses whole embryos without the isolation of pure nuclei, our protocol may not be suitable for biochemical experiments such as western blotting, proteomics, etc.

## Materials and Methods

The main experimental steps of *in situ* NuMat preparation have been outlined in the form of a workflow in Figure 1. A detailed step by step protocol has been submitted as Supplementary document 1.

### *Drosophila* embryo/tissue collection, fixation and permeabilization

*Drosophila* embryos (0-2 hr old) or of desired developmental age, were collected on a grape juice-agar collection plate. The collected embryos were dechorionated with 50% sodium hypochlorite and washed thoroughly with running tap water. The dechorionated embryos were then devitellinised and fixed simultaneously in a mix of 4% formaldehyde in Phosphate buffered saline (PBS) and heptane (in 1:1 v/v ratio). The embryos were vigorously shaken in the fixative:heptane mix for 20 mins at room temperature (RT). The aqueous layer at the bottom, was removed with a pipette and the embryos were further devitellinised using a mixture of 1:1 (v/v) ice-cold methanol:heptane. The tube was shaken vigorously until the embryos start to settle at the bottom of the tubes. Embryos that are devitellinised completely sink to the bottom, while the damaged ones and the ones that have vitelline membrane still attached, remain floating. This step was repeated several times. The devitellinised embryos were then equilibrated in aqueous media with several washes in PBS + 0.1% Triton-X-100 (PBT). Some embryos were reserved to serve as unextracted controls while the rest were used for NuMat preparation.

For *in situ* NuMat preparation from salivary glands or imaginal discs, the desired tissue was dissected out from 3^rd^ instar larvae and washed in PBS. The tissue was then fixed with 4% formaldehyde in PBT for 20 mins at RT. Fixative was removed by washing thrice with PBT. For each wash PBT was added to the tube and the tissue was allowed to settle with gravity following which the PBT was removed.

Efficient dechorionation and devitellinisation of embryos is a critical step of the protocol. Improper removal of chorion makes the embryo impermeable to most of the treatments thus compromising NuMat preparation. Fixation is another important step for *in situ* NuMat preparation. Over-fixation may lead to artefactual attachment of molecules to NuMat whereas under-fixation may cause extraction of genuine components. Here, we use 4% formaldehyde for fixation but alternative fixatives such as paraformaldehyde may be used, in which case the time required for fixation must be empirically determined. We have also isolated NuMat without fixation and observe that the nuclear shape, size and architecture as revealed by Lamin Dm0 immuno-staining, remains intact. Thus, the decision to fix or not to fix would depend on how labile the queried component is. However, coagulating fixatives such as ethanol or long-range fixatives such as DSG (Disuccinimidyl glutarate), EGS (Ethylene glycol bis(succinimidyl succinate)) etc. should be avoided because they may cause artefacts.

### *In situ* NuMat preparation

To prepare NuMat, fixed/unfixed embryos/tissue were stabilized by incubating for 20 mins at 37°C in PBT. This step stabilizes the nuclear architecture and facilitates the isolation of NuMat with comparatively intact composition as compared to unstabilized embryos/tissue. After stabilization, the embryos/tissue were extracted sequentially with salt and non-ionic detergent. Extraction was carried out first in 0.4M NaCl followed by 2M NaCl along with 0.5% Triton-X-100. During this step, most of the nucleoplasmic proteins are extracted. The high salt treatment removes majority of histones leading to unpackaging of DNA. The loosened DNA protrudes out from the nuclear margin and appears as a halo when stained with DNA dyes like DAPI. Salt extraction was followed by washes in PBT and then the DNA was removed by extensive digestion with DNase I. This treatment removes most of the DNA and chromatin associated components. The embryos/tissue were finally washed with PBT. These embryos/tissue containing *in situ* NuMat were further processed for immuno-staining, immuno-FISH or transmission electron microscopy (TEM) using standard protocols to visualize the queried component of nuclear architecture.

## Acknowledgements

The authors thank RKM lab members for helpful discussions. We thank Dr. Nandini Rangaraj and N.R. Chakravarthi for the help with confocal microscopy.

## References

1. Berezney, R. & Coffey, D. S. Identification of a nuclear protein matrix. Biochem. Biophys. Res. Commun. 60, 1410–1417 (1974).

2. Berezney, R. & Coffey, D. Nuclear protein matrix: association with newly synthesized DNA. Science (80-.). 189, 291–293 (1975).

3. Jackson, D. A. & Cook, P. R. Transcription occurs at a nucleoskeleton. EMBO J. 4, 919–925 (1985).

4. Mullenders, L. H. F., van Leeuwen, A. C. K. v., van Zeeland, A. A. & Natarajan, A. T. Nuclear matrix associated DNA is preferentially repaired in normal human fibroblasts, exposed to a low dose of ultraviolet light but not in Cockayne’s syndrome fibroblasts. Nucleic Acids Res. 16, 10607–10622 (1988).

5. Zeitlin, S., Parent, A., Silverstein, S. & Efstratiadis, A. Pre-mRNA splicing and the nuclear matrix. Mol Cell Biol 7, 111–120 (1987).

6. Reyes, J. C., Muchardt, C. & Yaniv, M. Components of the human SWI/SNF complex are enriched in active chromatin and are associated with the nuclear matrix. J. Cell Biol. 137, 263–274 (1997).

7. Pederson, T. Half a century of ‘the nuclear matrix’. Mol. Biol. Cell 11, 799–805 (2000).

8. He, D. C., Nickerson, J. a & Penman, S. Core filaments of the nuclear matrix. J. Cell Biol. 110, 569–80 (1990).

9. Jackson, D. a & Cook, P. R. The Structural Basis of Nuclear Function. International review of cytology 162A, 125–149 (1996).

10. Farrell, J. A. & O’Farrell, P. H. From egg to gastrula: How the cell cycle is remodeled during the drosophila mid-blastula transition. Annu. Rev. Genet. 48, 269–294 (2014).

11. Pathak, R. U., Rangaraj, N., Kallappagoudar, S., Mishra, K. & Mishra, R. K. Boundary Element-Associated Factor 32B Connects Chromatin Domains to the Nuclear Matrix. Mol. Cell. Biol. 27, 4796–4806 (2007).

12. Spradling, A. C. et al. Gene disruptions using P transposable elements: An integral component of the Drosophila genome project. Proc. Natl. Acad. Sci. U. S. A. 92, 10824–10830 (1995).

13. Spradling, A. C. et al. The Berkeley Drosophila Genome Project gene disruption project: Single P-element insertions mutating 25% of vital Drosophila genes. Genetics 153, 135–177 (1999).

14. Moore, H. M. & Vartiainen, M. K. F-actin organizes the nucleus. Nat. Cell Biol. 19, 1386–1388 (2017).

15. Fey, E. G. & Penman, S. Nuclear matrix proteins reflect cell type of origin in cultured human cells. Proc. Natl. Acad. Sci. U. S. A. 85, 121–125 (1988).

16. Jackson, D. A. Spatial epigenetics: Linking nuclear structure and function in higher eukaryotes. Essays Biochem. 48, 25–43 (2010).

17. Pathak, R. U. et al. AAGAG repeat RNA is an essential component of nuclear matrix in Drosophila. RNA Biol. 10, 564–71 (2013).

18. Capco, D. G., Wan, K. M. & Penman, S. The nuclear matrix: Three-dimensional architecture and protein composition. Cell 29, 847–858 (1982).

19. Wilson, R. H. C., Hesketh, E. L. & Coverley, D. The Nuclear Matrix: Fractionation Techniques and Analysis. Cold Spring Harb. Protoc. 2016, pdb.top074518(2016).

20. Breslin, S. & O’Driscoll, L. Three-dimensional cell culture: The missing link in drug discovery. Drug Discov. Today 18, 240–249 (2013).

21. Picollet-D’hahan, N. et al. A 3D Toolbox to Enhance Physiological Relevance of Human Tissue Models. Trends Biotechnol. 34, 757–769 (2016).

22. Kaufmann, S. H., Coffey, D. S. & Shaper, J. H. Considerations in the isolation of rat liver nuclear matrix, nuclear envelope, and pore complex lamina. Exp. Cell Res. 132, 105–123 (1981).

23. Kallappagoudar, S., Varma, P., Pathak, R. U., Senthilkumar, R. & Mishra, R. K. Nuclear Matrix Proteome Analysis of Drosophila melanogaster. Mol. Cell. Proteomics 9, 2005–2018 (2010).

24. Varma, P. & Mishra, R. K. Dynamics of nuclear matrix proteome during embryonic development in Drosophila melanogaster. J. Biosci. 36, 439–459 (2011).

25. Mamillapalli, A., Pathak, R. U., Garapati, H. S. & Mishra, R. K. Transposable Element ‘roo’ Attaches to Nuclear Matrix of the Drosophila melanogaster Transposable element ‘ roo ‘ attaches to nuclear matrix of the Drosophila melanogaster. 13, 1–27 (2013).

26. Pathak, R. U., Srinivasan, A. & Mishra, R. K. Genome-wide mapping of matrix attachment regions in Drosophila melanogaster. BMC Genomics 15, 1022(2014).

